# Polycomb suppresses a female gene regulatory network to ensure testicular differentiation

**DOI:** 10.1101/2021.01.19.427322

**Authors:** So Maezawa, Masashi Yukawa, Kazuteru Hasegawa, Ryo Sugiyama, Mengwen Hu, Miguel Vidal, Haruhiko Koseki, Artem Barski, Tony DeFalco, Satoshi H. Namekawa

## Abstract

Gonadal sex determination is controlled by the support cells of testes and ovaries. In testes, the epigenetic mechanism that maintains cellular memory to suppress female sexual differentiation remains unknown. Here, we show that Polycomb suppresses a female gene regulatory network in Sertoli cells, the specific support cells for postnatal testes. Through genetic ablation, we removed Polycomb repressive complex 1 (PRC1) from embryonic Sertoli cells after sex determination. PRC1-depleted postnatal Sertoli cells exhibited defective proliferation and cell death, leading to the degeneration of adult testes. In adult Sertoli cells, PRC1 suppressed the specific, critical genes required for granulosa cells, the support cells of ovaries, thereby inactivating the female gene regulatory network. The underlying chromatin of female genes was coated with Polycomb-mediated repressive modifications: PRC1-mediated H2AK119ub and PRC2-mediated H3K27me3. Taken together, we identify a critical mechanism centered on Polycomb that maintains the male fate in adult testes.

## Introduction

In mammals, gonadal sex determination takes place in the bipotential somatic cell precursors of male Sertoli cells and female granulosa cells in embryos (Capel, 2017; Swain & Lovell-Badge, 1999; Wilhelm, Palmer, & Koopman, 2007). Sertoli cells are the first somatic cells to differentiate in the XY gonad. In testes, Sertoli cells function as a regulatory hub for both differentiation and survival of germ cells, thereby determining male sexual fate (Svingen & Koopman, 2013). The mechanisms maintaining the male cellular identity of Sertoli cells are fundamental for adult testicular functions, including spermatogenesis and hormone production.

At the time of sex determination in embryos, the commitment to the male fate is triggered by the expression of the Y-linked *Sry* gene, and, subsequently, the female fate is suppressed (Capel, 2017; Swain & Lovell-Badge, 1999; Wilhelm et al., 2007). Distinct gene regulatory networks promote the male or female fate and are regulated by strong feedback loops that antagonize each other, canalizing one fate from the other (Capel, 2017). Sexual fate is interchangeable even after the initial commitment to Sertoli cells or granulosa cells with the removal of specific, critical transcription factors. The loss of *Dmrt1* in Sertoli cells leads to derepression of *Foxl2*, a master regulator of granulosa cell fate, and transdifferentiation of cell fate from Sertoli to granulosa cells (Matson et al., 2011; Matson & Zarkower, 2012). On the other hand, the loss of *Foxl2* in granulosa cells leads to derepression of the male gene network and transdifferentiation of cell fate from granulosa to Sertoli cells (Uhlenhaut et al., 2009). These findings, together with follow-up studies (Li et al., 2017; Lindeman et al., 2015; Minkina et al., 2014; Nicol et al., 2018; Zhao, Svingen, Ng, & Koopman, 2015), suggest that active repression of the alternate sexual fate is important for both testicular and ovarian function, even in adult life.

Epigenetic silencing mechanisms serve as molecular switches for the sex determination of bipotential somatic cell precursors. The deletion of a Polycomb gene *Cbx2* results in male to female reversal (Katoh-Fukui et al., 1998), which is mediated through suppression of genes required for the female fate (Garcia-Moreno et al., 2019). Additionally, regulation of H3K9 methylation is important for the male sexual fate (Kuroki et al., 2013). Although these studies highlight the importance of epigenetic mechanisms for initial sex determination, the epigenetic mechanisms by which male cellular identity is maintained through cell divisions and the proliferation of Sertoli cells remain to be determined.

Polycomb proteins suppress non-lineage-specific genes and define cellular identities of each lineage in stem cells and in development (Aloia, Di Stefano, & Di Croce, 2013; Ringrose & Paro, 2007; Simon & Kingston, 2013). In this study, we show that Polycomb suppresses the female gene regulatory network in postnatal Sertoli cells, thereby promoting the male gene regulatory network to ensure male cell fate. We generated loss-of-function mouse models of Polycomb repressive complex 1 (PRC1) in Sertoli cells after initial sex determination. We show that PRC1 is required for the proliferation of Sertoli cells, as well as the suppression of non-lineage-specific genes and the female gene regulatory network in Sertoli cells. Taken together, we identify a critical mechanism centered on Polycomb that maintains male fate in adult testes.

## Results

### PRC1 in Sertoli cells is required for the maintenance of spermatogenesis

In postnatal Sertoli cells (which are detected by the Sertoli cell marker GATA4), RNF2 is highly expressed and the RNF2-mediated epigenetic mark H2AK119ub is abundant (Figure 1A), which suggests PRC1 functions in these cells. To determine the function of PRC1, we generated a PRC1 loss-of-function mouse model by removing two redundant catalytic subunits, RNF2 and RING1 (Endoh et al., 2012). We generated a conditional deletion of *Rnf2* (*Rnf2*cKO) using *Amh*-Cre, which is expressed specifically in Sertoli cells after embryonic day 14.5 (E14.5) (Holdcraft & Braun, 2004), in a background of *Ring1*-knockout (KO) mice (*Amh*-Cre; *Rnf2*cKO; *Ring1*-KO: termed PRC1^*Amh*-Cre^cKO: PRC1AcKO). Although RNF2 appears to be the most active component in the heterodimeric E3 ligases of PRC1, the RNF2 paralog RING1 can partially compensate for the loss of RNF2 (Endoh et al., 2012). Therefore, we made a conditional deletion of *Rnf2* in a background of *Ring1*-KO mice, which are viable and do not have fertility defects (del Mar Lorente et al., 2000). This strategy enabled us to define the function of RNF2 without compensation from RING1, while also representing a “complete” loss-of-function of PRC1 as shown in testicular germ cells (Maezawa et al., 2017) and in other biological contexts (Endoh et al., 2012; Posfai et al., 2012; Yokobayashi et al., 2013). Since PRC1 has various components, including CBX2 (Gao et al., 2012), this strategy allows us to determine the global function of PRC1. At the same time, the use of *Amh*-Cre allowed us to test the function of PRC1, specifically after the completion of sex determination at E12.5.

**Figure 1.**
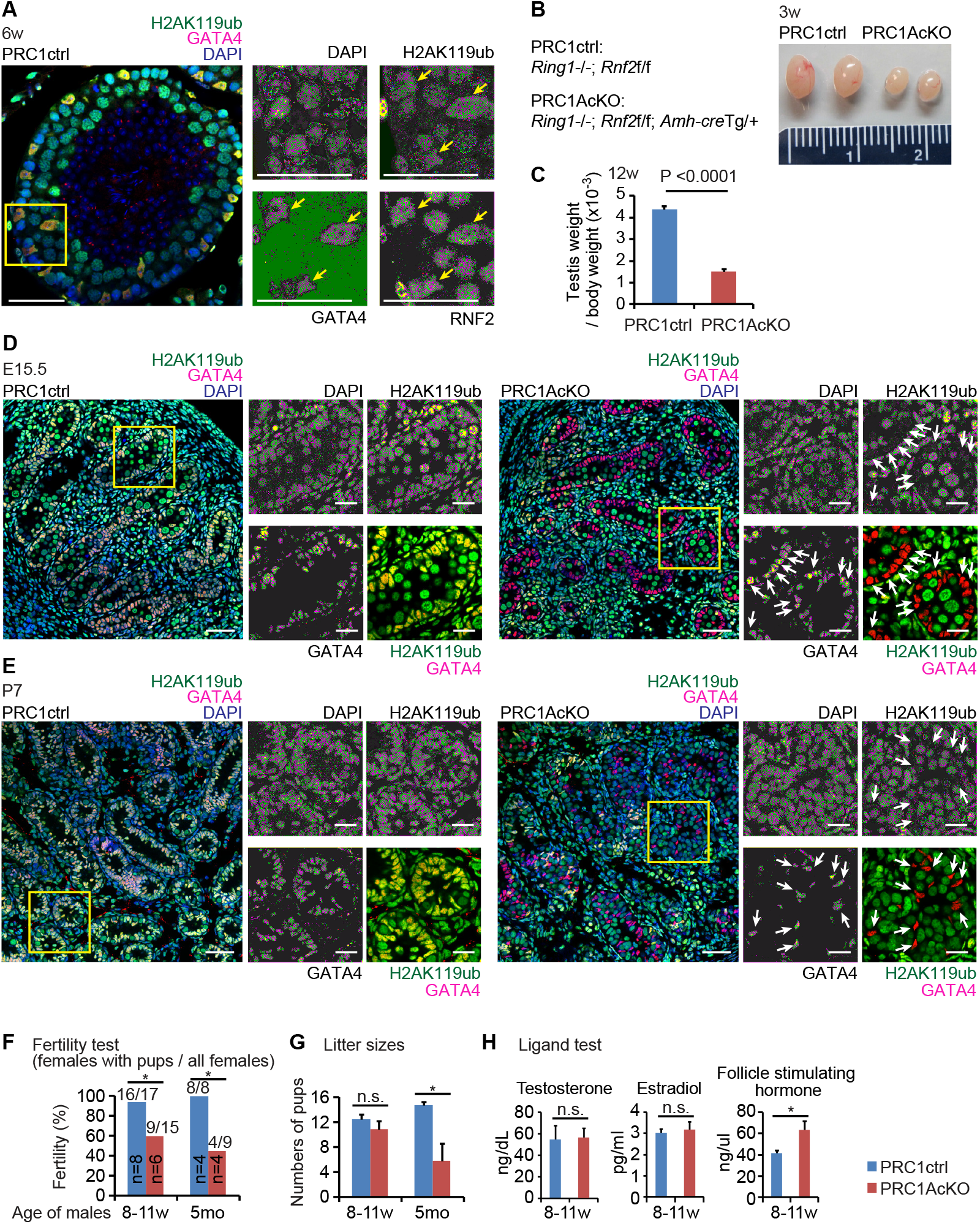
Deletion of PRC1 in Sertoli cells. (**A**) RNF2 and RNF2-mediated H2AK119ub localized at GATA4-positive Sertoli cells (yellow arrows) in a testicular section at 6 weeks of age. Regions bordered by yellow squares are magnified in the right panels. Bars in the large panels: 50 μm. Bars in the magnified panels: 20 μm. (**B**) Genotypes and photographs of testes at 3 weeks of age. Measurement scale in the panel: 2 cm. (**C**) Testicular weight/body weight ratio (× 10^−3^) at 12 weeks of age. *P* < 0.0001, unpaired t-test. (**D, E**) Localization of H2AK119ub and GATA4 in PRC1ctrl and PRC1AcKO at embryonic day 15.5 (**D**) and 1 week of age (**E**). Regions bordered by yellow squares are magnified in the right panels. Bars in the large panels: 50 μm. Bars in the magnified panels: 20 μm. H2AK119ub^-^ Sertoli cells in mutants are shown with white arrows. (**F**) The fertility of PRC1AcKO males, at 8-11 weeks of age and 5 months of age, were tested via crosses with CD1 wild-type females. Numbers of males tested are shown within the bars, and numbers of females with pups and all females are shown above the bars. *\*P* < 0.05, Fisher’s exact test. (**G**) Litter sizes of breeding tests at 8-11 weeks of age and 5 months of age. *\*P* < 0.05, Welch’s t-test. (**H**) Ligand hormone tests at 8-11 weeks of age testes. *\*P* < 0.05, Welch’s t-test. n.s., not significant.

PRC1AcKO males have smaller testes compared with littermate controls that harbored floxed alleles for *Rnf2* on a *Ring1*-KO background without *Amh*-Cre (termed PRC1 control: PRC1ctrl: Figure 1B and C). We confirmed efficient *Amh*-Cre-mediated recombination by observing depletion of the RNF2-mediated mark, H2AK119ub, in GATA4^+^ Sertoli cells of PRC1AcKO testes at E15.5 (> 95 % efficiency: Figure 1D) and at postnatal day 7 (P7: Figure 1E). In 6 week-old PRC1AcKO testes, while the tubules with H2AK119ub^+^ Sertoli cells (escaped Cre-mediated deletion) appear to have normal morphology, we frequently observed disorganization of testicular tubules that contain H2AK119ub^-^ Sertoli cells (underwent Cre-mediated deletion: arrowheads, Figure 1- figure supplement 1), suggesting a critical function of PRC1 in Sertoli cells in the organization of testicular tubules. This mosaic pattern is presumably due to incomplete Cre-mediated recombination, as the Sertoli cells that escaped Cre-mediated recombination might have repopulated the testes. Consistent with this interpretation, PRC1AcKO males are subfertile, and 9 out of 15 wild-type females mated with 8-11-week old PRC1AcKO males gave birth (Figure 1F) at comparable litter sizes (Figure 1G). Interestingly, fecundity decreased in aged PRC1AcKO males (5 months old), as litter sizes were smaller as compared to controls (Figure 1G). To further examine the phenotype, we measured the blood levels of three hormones critical for testicular homeostasis: although the levels of testosterone and estradiol were comparable between cKO and control mice, follicle-stimulating hormone was increased in mutants (Figure 1H). As follicle stimulating hormone levels are regulated by a feedback mechanism involving Sertoli cells (Oduwole, Peltoketo, & Huhtaniemi, 2018), we infer that the dysfunction of Sertoli cells and testicular degeneration caused increased follicle stimulating hormone levels to recover Sertoli cell function.

To determine the cause of testicular degeneration, we next examined the proliferation of Sertoli cells. In normal mouse development, Sertoli cells proliferate in fetal and neonatal testes until approximately 2 weeks after birth, and the number of Sertoli cells in adult testes determines both testis size and daily sperm production (Sharpe, McKinnell, Kivlin, & Fisher, 2003). In P7 control testes, GATA4^+^ Sertoli cells occasionally co-expressed an S phase marker, PCNA, and another marker of active cell cycle, Ki67 (Figure 2A), consistent with the active proliferation of Sertoli cells. However, in PRC1AcKO testes, GATA4^+^ Sertoli cells were largely devoid of PCNA and Ki67 (Figure 2A), suggesting impaired proliferation of PRC1AcKO Sertoli cells. Next, we independently confirmed the proliferation phenotype by using EdU labeling of actively proliferating cells. EdU was abdominally administrated to P7 mice, and testicular sections were examined the following day. While GATA4^+^ Sertoli cells were occasionally EdU^+^ (approximately one-fouth) in control testes, GATA4^+^ Sertoli cells were devoid of EdU signal in PRC1AcKO testes (Figure 2B). Additional labeling of H2AK119ub confirmed the loss of PRC1 function in GATA4^+^ Sertoli cells from PRC1AcKO testes (Figure 2B). From these results, we conclude that the loss of PRC1 disrupts the proliferation of Sertoli cells. This is in contrast with PRC1’s function in testicular germ cells, in which the loss of PRC1 does not affect proliferation but instead causes apoptotic cell death (Maezawa et al., 2017). This difference suggests a unique function of PRC1 in Sertoli cells that is distinct from its function in germ cells.

**Figure 2.**
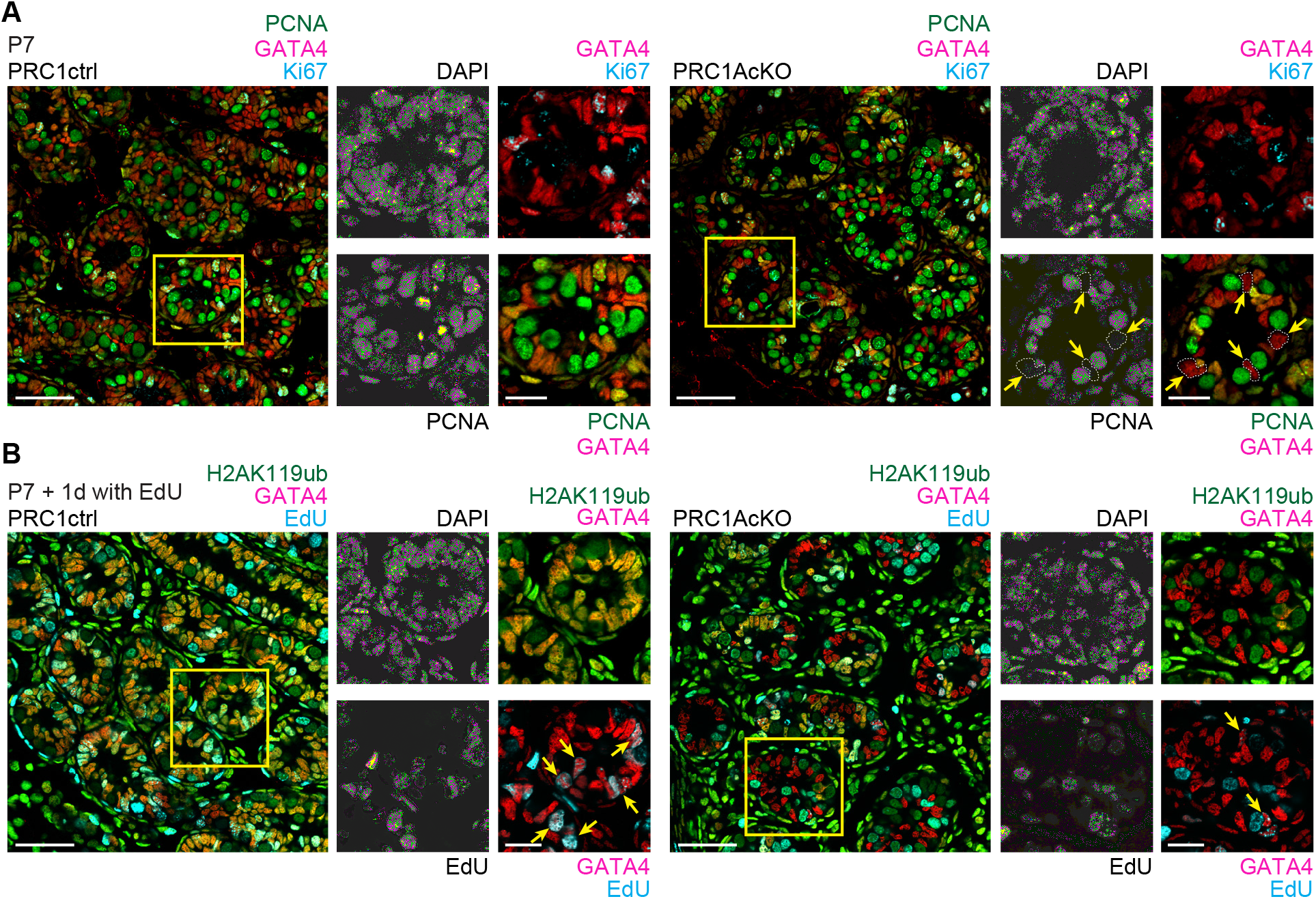
PRC1 is required for proliferation of Sertoli cells. (**A**) PCNA and Ki67 were not detected in GATA4-positive Sertoli cells (arrows) in a PRC1AcKO testicular section at 6 weeks of age, while PCNA and Ki67 were present in PRC1ctrl Sertoli cells. (**B**) Testicular sections of the indicated genotypes 1 day following the injection of EdU into males at 1 week of age. The presence of EdU-positive Sertoli cells (arrows) was decreased in PRC1AcKO testes. Regions bordered by yellow squares are magnified in the right panels. Bars in the large panels: 50 μm. Bars in the magnified panels: 20 μm.

### In Sertoli cells, PRC1 suppresses genes required for granulosa cells

We next sought to determine the genes regulated by PRC1 in Sertoli cells. In PRC1AcKO testes, some Sertoli cells escaped *Amh*-Cre-mediated recombination. Thus, it was difficult to specifically isolate Sertoli cells that underwent PRC1 depletion. To precisely determine the function of PRC1 in gene regulation in Sertoli cells, we used an alternative strategy: we isolated Sertoli cells from a mouse line in which conditional deletion of PRC1 can be induced by tamoxifen-inducible Cre-mediated recombination under the control of the endogenous ROSA26 promoter (*ROSA26*-Cre^ERT^; *Rnf2*^floxed/floxed^; *Ring1*-KO: termed PRC1^ROSA26-CreERT^cKO: PRC1RcKO). After isolating Sertoli cells from P7 testes, we cultured Sertoli cells for 4 days in the presence of 4-hydroxytamoxifen (4-OHT), and performed RNA-sequencing (RNA-seq: Figure 3A). As a control, we isolated Sertoli cells from control mice (*Rnf2*^floxed/floxed^; *Ring1*-KO) and cultured them in the same 4-OHT conditions. We performed RNA-seq for two independent biological replicates and confirmed reproducibility between biological replicates (Figure 3- figure supplement 1A).

**Figure 3.**
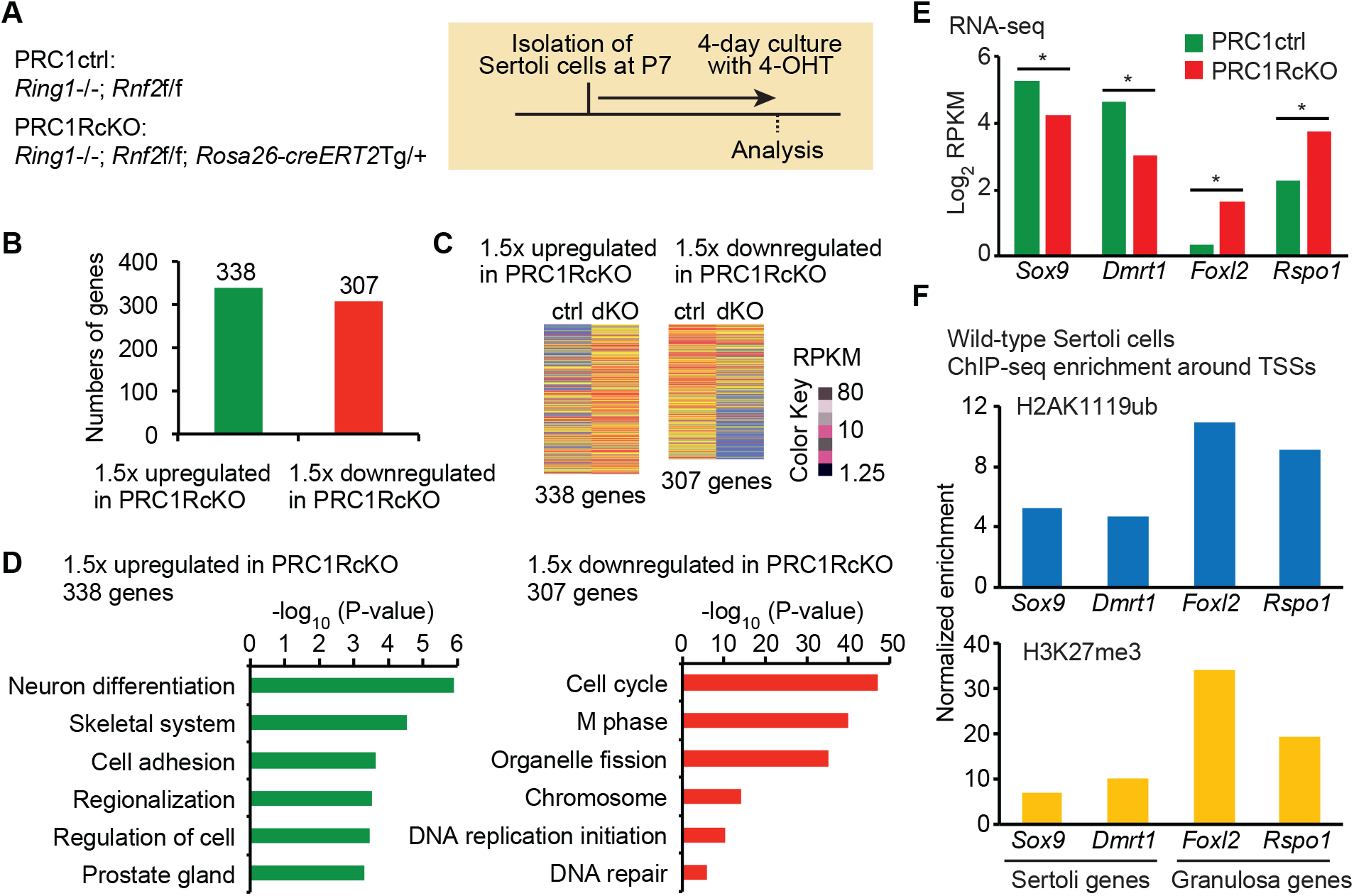
In Sertoli cells, PRC1 suppresses genes required for granulosa cells. (**A**) Genotypes and experiment schematic. Sertoli cells were isolated from P7 testes and cultured for 4 days in the presence of 4-OHT prior to RNA-seq analyses. (**B**) The numbers of differentially expressed genes detected by RNA-seq (≥1.5-fold change, *P*_*adj*_ < 0.05) in Sertoli cells (Two biological replicates) between PRC1ctrl and PRC1RcKO. (**C**)Heatmaps showing gene expression patterns for upregulated (left) and down-regulated (right) genes in Sertoli cells. (**D**) GO term analyses. (**E**) Expression levels for representative Sertoli and granulosa genes. (**F, G**) H2AK119ub and H3K27me3 ChIP-seq enrichment around the TSSs of representative Sertoli and granulosa genes.

Our RNA-seq analyses demonstrated that 338 genes were upregulated in PRC1RcKO Sertoli cells as compared to controls, while 307 genes were downregulated in PRC1RcKO Sertoli cells (Figure 3A and B). Gene ontology (GO) analysis showed that upregulated genes were enriched for functions in neural differentiation, skeletal system, and cell adhesion (Figure 3D). These categories suggest that PRC1 suppressed expression of non-lineage-specific genes in Sertoli cells. On the other hand, GO analysis revealed that downregulated genes were enriched for functions in the cell cycle and M phase (Figure 3D). This result is in accord with the cell cycle arrest we found in PRC1AcKO Sertoli cells (Figure 2).

Since we anticipated suppression of genes required for granulosa cells by PRC1 in Sertoli cells, we next investigated the expression level of key genes required for granulosa cells. In PRC1RcKO Sertoli cells, genes required for female sexual development were upregulated: these genes included *Rspo1*, an activator of the Wnt pathway (Chassot et al., 2008), and *Foxl2*, a key transcription factor for granulosa cells (Schmidt et al., 2004) (Figure 3E). Importantly, these female genes suppress the male fate, and the loss of these genes leads to female-to-male sex reversal (Ottolenghi et al., 2007; Parma et al., 2006; Schmidt et al., 2004). Consistent with the antagonistic function of these female genes with the male pathway, key sex determination genes for the male pathway were downregulated in PRC1RcKO Sertoli cells: these genes included *Sox9*, an evolutionarily conserved gene for sex determination, which directs the male pathway downstream of *Sry* (Vidal, Chaboissier, de Rooij, & Schedl, 2001),and *Dmrt1* (Raymond, Murphy, O’Sullivan, Bardwell, & Zarkower, 2000; Raymond et al., 1998), male-determining signalling (Figure 3E).

To determine whether the suppression of female genes was directly regulated by PRC1, we performed chromatin immunoprecipitation sequencing (ChIP-seq) of PRC1-mediated H2AK119ub in isolated wild-type Sertoli cells. We further performed ChIP-seq of H3K27me3 in Sertoli cells since PRC2-mediated H3K27me3 is regulated by PRC1 and its mediated mark H2A119ub (Blackledge et al., 2014; Cooper et al., 2014). We performed ChIP-seq for two independent biological replicates and confirmed the reproducibility between biological replicates (Figure 3- figure supplement 1B). We confirmed the enrichment of H2AK119ub and H3K27me3 around transcription start sites (TSSs) of *Rspo1* and *Foxl2* (Figure 3F). Compared to the enrichment of H2AK119ub on these female genes, enrichment of H2AK119ub was relatively low on the TSSs of the male genes, *Sox9* and *Dmrt1*. Therefore, we conclude that PRC1 directly binds and suppresses *Rspo1* and *Foxl2*.

Track views of these ChIP-seq data show that all three marks were enriched at *Foxl2* and *Rspo1* loci (Figure 4A). Furthermore, enrichment of these three marks was found in other loci such as *Foxf2*, the mutation of which appears in patients with disorders of sex development (Jochumsen et al., 2008), and *Hoxd13*, which is involved in female reproductive tract development (Du & Taylor, 2015). These results suggest that PRC1 works with PRC2 to suppress female genes as well as developmental regulators such as *Fox* and *Hox* genes, which are the classical targets of Polycomb-mediated gene repression (Lee et al., 2006).

**Figure 4.**
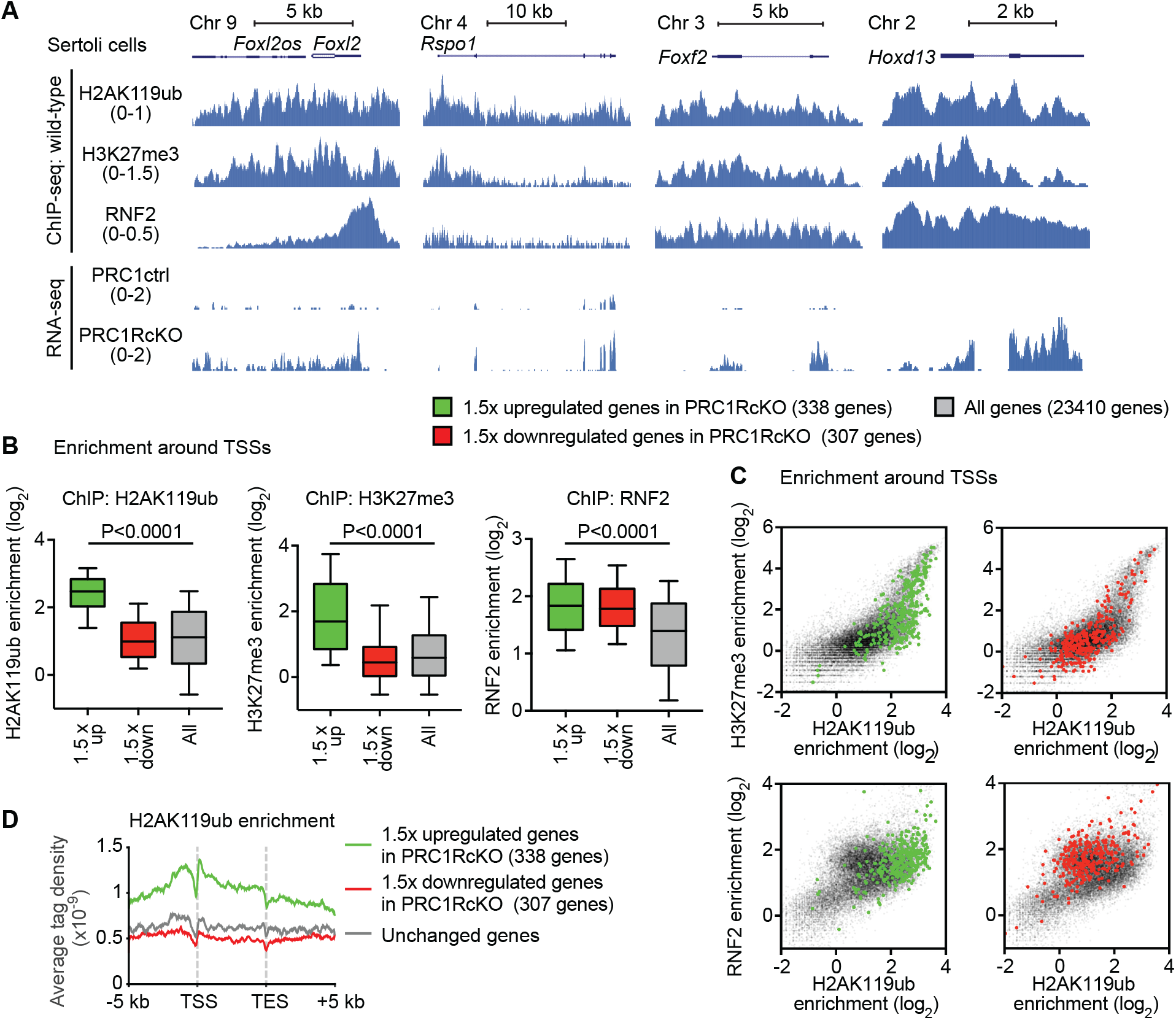
In Sertoli cells, Polycomb-mediated marks are enriched on genes required for granulosa cells. (**A**) Genome track views of representative genes in the female gene regulatory network. ChIP-seq enrichment in wild-type Sertoli cells is shown (top); RNA-seq peaks in PRC1ctrl and PRC1RcKO Sertoli cells are shown (bottom). (**B**) Box-and-whisker plots showing distributions of enrichment for ChIP-seq data. Central bars represent medians, the boxes encompass 50% of the data points, and the whiskers indicate 90% of the data points. *P*, Mann-Whitney U tests. (**C**) Scatter plots showing ChIP-seq enrichment (±2 kb around TSSs) of indicated modifications on genes upregulated (left panels) and down-regulated (right panels) in Sertoli cells. The distribution of all genes is shown with gray dots. (**D**)Average tag densities of H2AK119ub ChIP-seq enrichment.

To determine the features of genome-wide gene repression mediated by Polycomb complexes, we analyzed the enrichment of H2AK119ub, H3K27me3, and RNF2 on the upregulated genes in PRC1RcKO Sertoli cells. H2AK119ub, H3K27me3, and RNF2 were all significantly enriched on the TSSs of upregulated genes in PRC1RcKO Sertoli cells as compared to all genes in the genome (Figure 4B). Additional enrichment analysis confirmed the co-enrichment of H2AK119ub and H3K27me3 (Figure 4C, upper panels) as well as co-enrichment of H2AK119ub and RNF2 (Figure 4C, lower panels) on upregulated genes in PRC1RcKO Sertoli cells. Furthermore, average tag density analysis confirmed that enrichment of H2AK119ub on upregulated genes in PRC1RcKO Sertoli cells occured on upstream regions, gene bodies, and downstream regions with the highest enrichment near TSSs (Figure 4D). We found a similar distribution of H3K27me3 around the gene bodies of upregulated genes in PRC1RcKO Sertoli cells (Figure 4- figure supplement 1). Together, these results confirmed the genome-wide, global functions of PRC1 in the direct regulation of gene repression in Sertoli cells.

### Polycomb globally inactivates the female gene regulatory network in postnatal Sertoli cells

Previous studies have suggested that sex determination is canalized by the interconnected, antagonistic network of genes both in males and in females that are controlled by feedback mechanisms (Capel, 2017). Since Polycomb is implicated in the maintenance of sex-specific gene regulatory networks through the PRC2-mediated mark H3K27me3 (Garcia-Moreno et al., 2019), we hypothesized that PRC1 globally inactivates the female gene regulatory network in postnatal Sertoli cells. Since male- and female-specific gene networks are regulated immediately after sex determination during fetal stages (Jameson et al., 2012), we reasoned that female gene network suppression in fetal Sertoli cells is maintained by PRC1 in postnatal Sertoli cells. To test this hypothesis, we examined the expression profiles of specifically expressed genes in E13.5 granulosa cells (Jameson et al., 2012) (Figure 5- figure supplement 1), termed “pre-granulosa genes” in postnatal Sertoli cells. We found that pre-granulosa genes were upregulated in PRC1RcKO postnatal Sertoli cells as compared to other genes (Figure 5A). On the other hand, specifically expressed genes in E13.5 Sertoli cells (Jameson et al., 2012) (Figure 5- figure supplement 1), termed “pre-Sertoli genes,” were downregulated in PRC1RcKO postnatal Sertoli cells as compared to other genes (Figure 5A). We further examined the correlation of each gene and found that pre-granulosa genes were highly correlated with upregulated genes in PRC1RcKO postnatal Sertoli cells, while pre-Sertoli genes were highly correlated with downregulated genes in PRC1RcKO postnatal Sertoli cells (Figure 5B). These results suggest that PRC1 maintains suppression of the female gene regulatory network, which is initiated at the time of sex determination and is maintained throughout the development of postnatal Sertoli cells.

**Figure 5.**
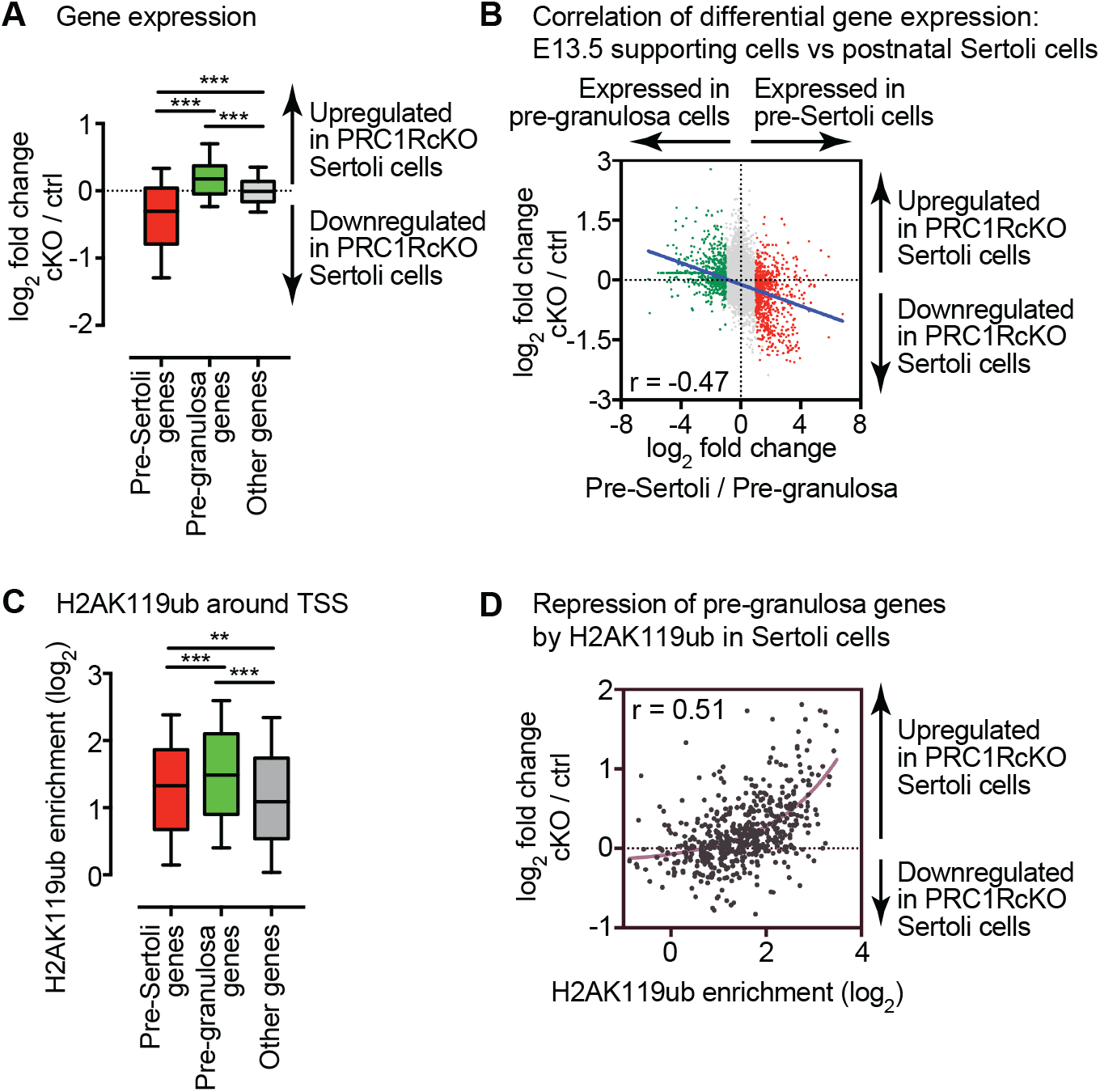
Polycomb inactivates the female gene regulatory network in Sertoli cells. Box-and-whisker plots showing distributions of RNA-seq data. Central bars represent medians, the boxes encompass 50% of the data points, and the whiskers indicate 90% of the data points. *\*\*\* P* < 0.0001, Mann-Whitney U tests. (**B**) Scatter plots showing the Pearson correlation between RNA-seq data for genes regulated in E13.5 support cells and P7 Sertoli cells. A linear trendline is shown in blue. (**C**) Box-and-whisker plots showing distributions of enrichment for H2AK119ub ChIP-seq data. Central bars represent medians, the boxes encompass 50% of the data points, and the whiskers indicate 90% of the data points. *\*\*\* P* < 0.0001, *\*\* P <* 0.005, Mann-Whitney U tests. (**D**) Scatter plots showing the Pearson correlation between ChIP-seq enrichment (±2 kb around TSSs) and gene expression in Sertoli cells. A linear trendline is shown in blue.

We next sought to determine whether PRC1 directly suppresses the female gene regulatory network in postnatal Sertoli cells. H2AK119ub is significantly enriched on TSSs of pre-granulosa genes compared to other genes and pre-Sertoli genes (Figure 5C). Among pre-granulosa genes, enrichment of H2AK119ub is positively correlated with upregulated genes in PRC1RcKO postnatal Sertoli cells (Figure 5D). We further identified the enrichment of H3K27me3 on TSSs of pre-granulosa genes in postnatal Sertoli cells (Figure 5- figure supplement 2A), and the correlation between H3K27me3 and upregulated genes on the pre-granulosa genes in PRC1RcKO postnatal Sertoli cells (Figure 5- figure supplement 2B). Together, we conclude that PRC1 globally inactivates the female gene regulatory network in postnatal Sertoli cells.

## Discussion

In this study, we demonstrated that PRC1 is required for proliferation of Sertoli cells and suppression of the non-lineage-specific gene expression program and the female gene regulatory network. Among these functions, we infer that suppression of the female gene regulatory network is the key mechanism to ensure male fate in addition to canonical functions of PRC1, which controls proliferation and suppression of non-lineage-specific gene expression programs. Although we did not observe complete infertility using *Amh*-Cre, presumably due to the repopulation of Sertoli cells that escaped Cre-mediated recombination, smaller testis size and abnormal tubule organization of cKO testes (Figure 1) suggest that PRC1 is critical for physiological functions of Sertoli cells. Below, we discuss two molecular aspects underlying these physiological phenotypes: control of proliferation and suppression of the female gene regulatory network.

The proliferation of Sertoli cells is a critical determinant of testicular functions because the number of Sertoli cells defines testicular functions (Sharpe et al., 2003). Rapid proliferation is a prominent feature of juvenile Sertoli cells. In general, Polycomb proteins are associated with the cell cycle checkpoint by directly suppressing the tumor suppressor locus *Cdkn2a*/*Ink4a*/*Arf* (Jacobs, Kieboom, Marino, DePinho, & van Lohuizen, 1999), which functions as a barrier to cancer transformation (Serrano et al., 1996). Therefore, enhanced Polycomb activity is a frequent feature of human tumors. We found that *Cdkn2a* was derepressed in PRC1RcKO Sertoli cells (approximately 2.5-fold upregulation: PRKM value for PRC1RcKO: 32.1 v.s. PRC1ctrl: 13.0). Therefore, our results suggest that PRC1 promotes the rapid proliferation of Sertoli cells by suppressing *Cdkn2a*. The function of PRC1 in Sertoli cell proliferation is distinct from Polycomb functions in testicular germ cells, where PRC1 depletion does not alter the proliferation of germ cells (Maezawa et al., 2017). This may be due to the fact that PRC1 and PRC2 are not required for suppression of *Cdkn2a* in germ cells (Maezawa et al., 2017; Mu, Starmer, Fedoriw, Yee, & Magnuson, 2014). The functional difference in testicular germ cells and Sertoli cells highlights the context-dependent functions of Polycomb proteins. However, a recent study revealed a novel activity by which PcGs can regulate cell proliferation through DNA replication independently of *Cdkn2a* (Piunti et al., 2014). Therefore, determining the detailed molecular mechanisms by which PRC1 controls the proliferation of Sertoli cells will be important for future studies.

Another critical function of PRC1 in Sertoli cells is the suppression of the female gene regulatory network. DMRT1 is a critical transcription factor that suppresses the expression of female genes (Matson et al., 2011). While more than 10-fold upregulation of the female genes (*Rspo1* and *Foxl2*) was observed for *Dmrt1* mutants (Matson et al., 2011), the degree of upregulation of these genes was modest in PRC1RcKO Sertoli cells (Figure 3). This finding, combined with genetic evidence, indicates that DMRT1 could be the direct regulator of suppression of female genes, while PRC1-mediated mechanisms could be a maintenance mechanism in response to the primary silencing mechanisms determined by DMRT1. Another possibility is compensation by PRC1-independent suppression mechanisms. While a portion of PRC2-mediated H3K27me3 is regulated by variant PRC1 (Blackledge et al., 2014; Cooper et al., 2014), another portion of H3K27me3, mediated by canonical PRC1 and PRC2 complexes, is not downstream of PRC1 (Laugesen, Hojfeldt, & Helin, 2019). These PRC2-mediated mechanisms or other silencing machinery may be responsible for the suppression of female genes.

The notion that Polycomb regulates the female gene regulatory network has been supported by other recent evidence. In bipotential precursor cells, genes involved in sex determination are marked with bivalent chromatin domains (Garcia-Moreno et al., 2019) that are prevalent in pluripotent stem cells and in germ cells (Bernstein et al., 2006; Hammoud et al., 2009; Lesch, Dokshin, Young, McCarrey, & Page, 2013; Maezawa, Hasegawa, et al., 2018; Maezawa, Yukawa, Alavattam, Barski, & Namekawa, 2018; Sin, Kartashov, Hasegawa, Barski, & Namekawa, 2015). Maintenance of the male fate was explained by the persistence of H3K27me3 on silent female genes in Sertoli cells (Garcia-Moreno et al., 2019). Consistent with PRC1’s function found in the current study, Polycomb-mediated silencing may globally suppress the female gene regulatory network. We found that H2AK119ub was largely associated with this group of female genes (Figure 5). Therefore, it would be interesting to speculate that antagonistic male and female networks can be directly coordinated by Polycomb protein functions, including the strong feedback mechanism underlying both networks. These possibilities raise several outstanding questions to be addressed in future studies. What are the functions of another Polycomb complex, PRC2, and of each Polycomb complex component, including CBX2, in postnatal Sertoli cells? What is the function of Polycomb in female granulosa cells, especially in the suppression of the male gene regulatory network? Does Polycomb underlie the feedback regulation of each network to define sexual identity? Our current study provides a foundation to explore these questions.

## Methods

### Animals

Generation of mutant *Ring1* and *Rnf2* floxed alleles were previously reported (Cales et al., 2008). *Amh-Cre* transgenic mice were purchased from The Jackson Laboratory (Stock No: 007915) (Holdcraft & Braun, 2004). *Rosa-Cre ERT* mice were purchased from The Jackson Laboratory (Stock No: 008463) (Ventura et al., 2007). A minimum of three independent mice were analyzed for each experiment. All of the animals were handled according to approved institutional animal care and use committee (IACUC) protocols (#IACUC2018-0040) of Cincinnati Children’s Hospital Medical Center. Fertility tests were performed with 6-weeks old CD1 female mice (purchased from Charles river). At least 2 female mice were bred with a male mouse for 2 weeks, and the fertility was evaluated by the ratio of pregnant to total female mice and number of pups.

### Ligand test

Blood samples were collected from C57BL/6N mice aged 8-11 weeks. Serum was separated immediately and stored at -20°C. Hormone assays, including testosterone, estradiol, and follicle stimulating hormone, were performed by the Center for Research in Reproduction at the University of Virginia.

### Sertoli cell isolation

Sertoli cells were isolated as previously described with minor modifications (Chang, Lee-Chang, Panneerdoss, MacLean, & Rao, 2011) and collected from C57BL/6N mice aged 6–8 days. Testes were collected in a 24-well plate in Dulbecco’s Modified Eagle Medium (DMEM) supplemented with GlutaMax (Thermo Fisher Scientific), non-essential amino acids (NEAA) (Thermo Fisher Scientific), and penicillin and streptomycin (Thermo Fisher Scientific). After removing the *tunica albuginea* membrane, testes were digested with collagenase (1 mg/ml) at 34°C for 20 min to remove interstitial cells, then centrifuged at 188×*g* for 5 min. Tubules were washed with medium and then digested with trypsin (2.5 mg/ml) at 34°C for 20 min to obtain a single-cell suspension. To remove KIT-positive spermatogonia, cells were washed with magnetic cell-sorting (MACS) buffer (PBS supplemented with 0.5% BSA and 5 mM EDTA) and incubated with CD117 (KIT) MicroBeads (Miltenyi Biotec) on ice for 20 min. Cells were separated by autoMACS Pro Separator (Miltenyi Biotec) with the program “possel.” Cells in the flow-through fraction were washed with MACS buffer and incubated with CD90.2 (THY1) MicroBeads (Miltenyi Biotec) on ice for 20 min to remove THY1-positive spermatogonia. Cells were separated by autoMACS Pro Separator (Miltenyi Biotec) with the program “posseld.” Cells in the flow-through fraction were washed and plated in a 6-well plate for 1 h in the medium supplemented with 10% fetal bovine serum, which promotes adhesion of Sertoli cells. Purity was confirmed by immunostaining.

For PRC1RcKO, cells were cultured for 4 days with 1 µM 4-OHT in Dulbecco’s Modified Eagle Medium (DMEM) supplemented with GlutaMax (Thermo Fisher Scientific), non-essential amino acids (NEAA) (Thermo Fisher Scientific), and penicillin and streptomycin (Thermo Fisher Scientific). The same medium was replaced 2 days after the initiation of the culture.

### Histological analysis and germ cell slide preparation

For the preparation of testicular paraffin blocks, testes were fixed with 4% paraformaldehyde (PFA) overnight at 4°C with gentle inverting. Testes were dehydrated and embedded in paraffin. For histological analysis, 7 µm-thick paraffin sections were deparaffinized and stained with hematoxylin and eosin. For immunofluorescence analysis of testicular sections, antigen retrieval was performed by boiling the slides in target retrieval solution (DAKO) for 10 min and letting the solution cool for 30 min. Sections were blocked with Blocking One Histo (Nacalai) for 1 h at room temperature and then incubated with primary antibodies overnight at 4°C. The resulting signals were detected by incubation with secondary antibodies conjugated to fluorophores (Thermo Fisher Scientific). Sections were counterstained with DAPI. Images were obtained via a laser scanning confocal microscope A1R (Nikon) and processed with NIS-Elements (Nikon) and ImageJ (National Institutes of Health) (Schneider, Rasband, & Eliceiri, 2012).

### ChIP-sequencing, RNA-sequencing, and data analysis

RNA-seq analyses were performed in the BioWardrobe Experiment Management System (Kartashov & Barski, 2015). Briefly, reads were aligned by STAR (version STAR_2.5.3a) with default arguments except --outFilterMultimapNmax 1 and --outFilterMismatchNmax 2. The -- outFilterMultimapNmax parameter was used to allow unique alignments only, and the -- outFilterMismatchNmax parameter was used to allow a maximum of 2 errors. NCBI RefSeq annotation from the mm10 UCSC genome browser was used, and canonical TSSs (1 TSS per gene) were analyzed. All reads from the resulting .bam files were split for related isoforms with respect to RefSeq annotation. Then, the EM algorithm was used to estimate the number of reads for each isoform. To detect differentially expressed genes between two biological samples, a read count output file was input to the DESeq2 package (version 1.16.1); then, the program functions DESeqDataSetFromMatrix and DESeq were used to compare each gene’s expression level between two biological samples. Differentially expressed genes were identified through binominal tests, thresholding Benjamini-Hochberg-adjusted P values to <0.01. To perform gene ontology analyses, the functional annotation clustering tool in DAVID (version 6.8) was used, and a background of all mouse genes was applied. Biological process term groups with a significance of P < 0.05 (modified Fisher’s exact test) were considered significant.

Cross-linking ChIP-seq with the ChIPmetation method (Schmidl, Rendeiro, Sheffield, & Bock, 2015) was performed for H2AK119ub, H3K27me3, and RNF2 as described previously (Schmidl et al., 2015). Data analysis for both ChIP-seq and RNA-seq was performed in the BioWardrobe Experiment Management System (https://github.com/Barski-lab/biowardrobe (Kartashov & Barski, 2015)). Briefly, reads were aligned to the mouse genome (mm10) with Bowtie (version 1.0.0 (Langmead, Trapnell, Pop, & Salzberg, 2009)), assigned to RefSeq genes (which have one annotation per gene) using the BioWardrobe algorithm, and displayed on a local mirror of the UCSC genome browser as coverage. Peaks of H2AK119ub-, H3K27me3- and RNF2-enrichment were identified using MACS2 (version 2.0.10.20130712 (Zhang et al., 2008)). Pearson correlations for the genome-wide enrichment of the peaks among ChIP-seq library replicates were analyzed using SeqMonk (Babraham Institute). Average tag density profiles were calculated around gene bodies, including 5-kb upstream and 5-kb downstream of the genes. Resulting graphs were smoothed in 200-bp windows. Enrichment levels for ChIP-seq experiments were calculated for 4-kb windows, promoter regions of genes (±2 kb surrounding TSSs), and enhancer regions. To normalize tag value read counts were multiplied by 1,000,000 and then divided by the total number of reads in each nucleotide position. The total amount of tag values in promoter or enhancer regions were calculated as enrichment. Microarray data was analyzed using the processed data (Jameson et al., 2012). Differentially expressed genes were identified through a p-value cutoff of 0.05, and a fold change cutoff of 2 for the comparison between E13.5 XX supporting cells and E13.5 XY supporting cells. Highly expressed genes in E13.5 XY supporting cells and in E13.5 XX supporting cells were termed as “Pre-sertoli genes” and “Pre-granulosa genes”, respectively. RNA-seq and ChIP-seq data reported in this paper have been deposited in GEO under accession code GSE167516.

## Acknowledgments

We thank David Zarkower and Vivian Bardwell for their help and discussion in the initial phase of this project, Katie Gerhardt for editing the manuscript, and the members of the Namekawa and Maezawa laboratories for discussion and helpful comments regarding the manuscript. This work was supported by Japan Society for the Promotion of Science Grant-in-Aid for Research Activity Start-up (19K21196), the Takeda Science Foundation (2019), and the Uehara Memorial Foundation Research Incentive Grant (2018) to SM, NIH Grants DP2GM119134 to AB, R35GM119458 to TD, and R01GM098605 and R01GM122776 to SHN.

---

**Figure 1- figure supplement 1.**
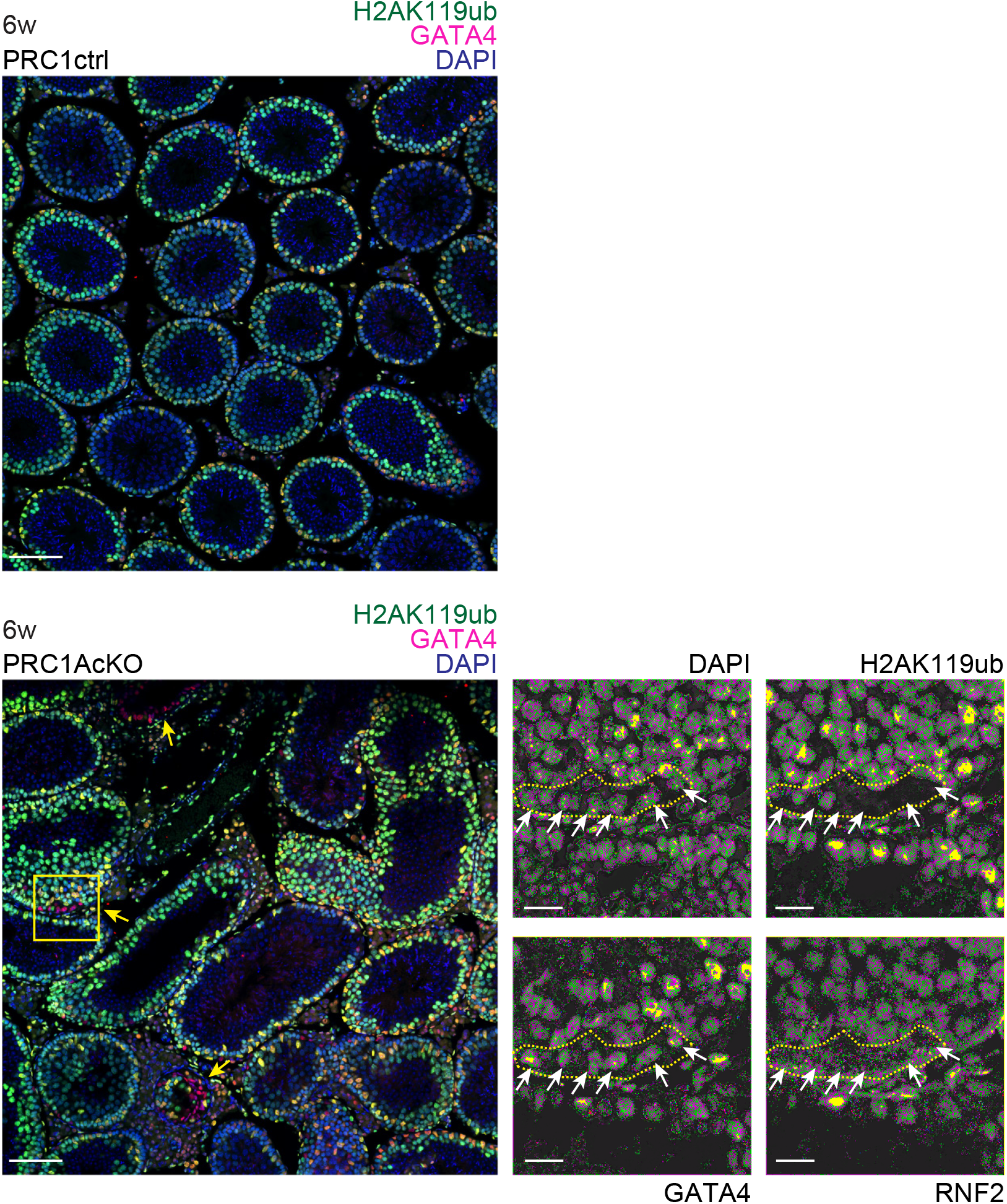
Deletion of PRC1 in Sertoli cells causes degeneration of testes at 6 weeks of age. Localization of H2AK119ub and GATA4 in PRC1ctrl and PRC1AcKO at 6 weeks of age. Regions bordered by yellow squares are magnified in the right panels. Bars in the large panels: 50 μm. Bars in the magnified panels: 20 μm. H2AK119ub^-^ Sertoli cells in mutants are shown with white arrows.

**Figure 3- figure supplement 1.**
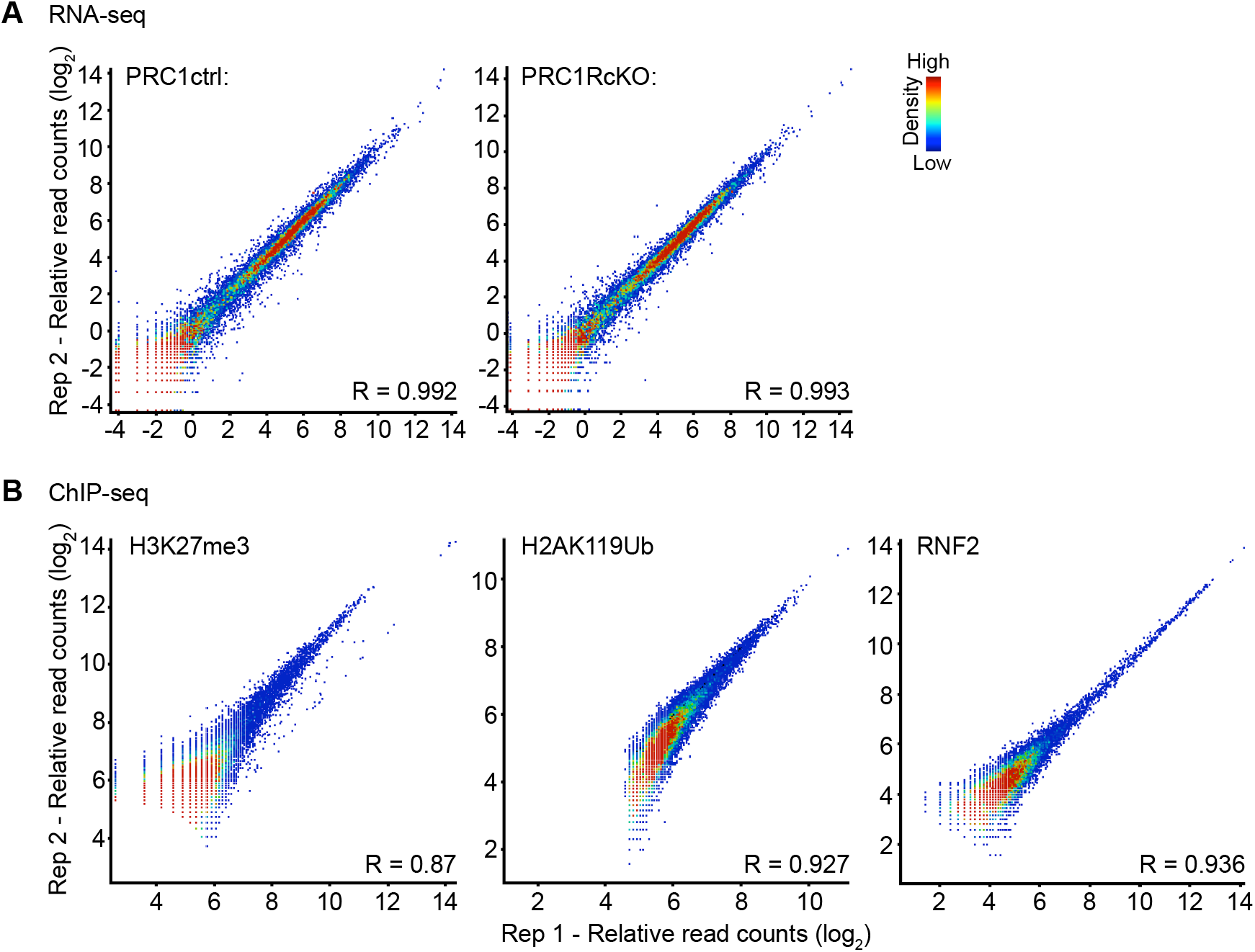
Biological replicates for RNA-seq and ChIP-seq data. (**A**) Scatter plots show the reproducibility of RNA-seq enrichment at individual peaks between biological replicates. (**B**) Scatter plots show the reproducibility of and ChIP-seq enrichment at individual peaks between biological replicates. Each peak was identified using MACS (P < 1×10^−5^). H3K27ac ChIP-seq enrichment levels are shown in log_2_ RPKM values. The color scale indicates RNA-seq or ChIP-seq peak density. Pearson correlation values (R) are shown.

**Figure 4- figure supplement 1.**
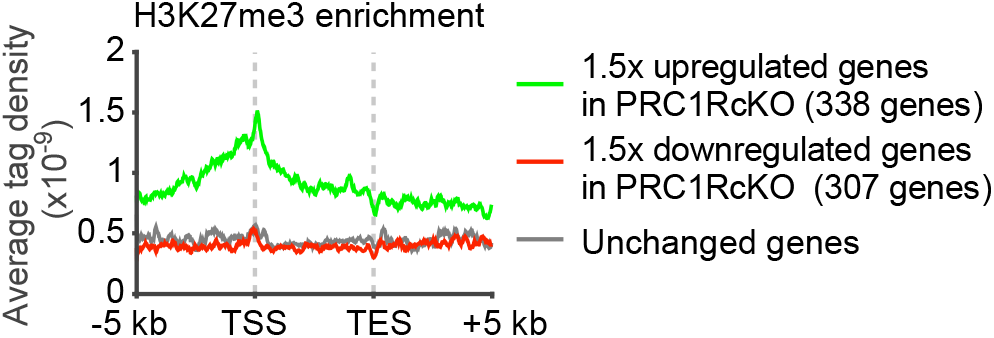
In Sertoli cells, H3K27me3 is enriched on genes required for granulosa cells. Average tag densities of H3K27me3 ChIP-seq enrichment on the groups of genes indicated in the panel.

**Figure 5- figure supplement 1.**
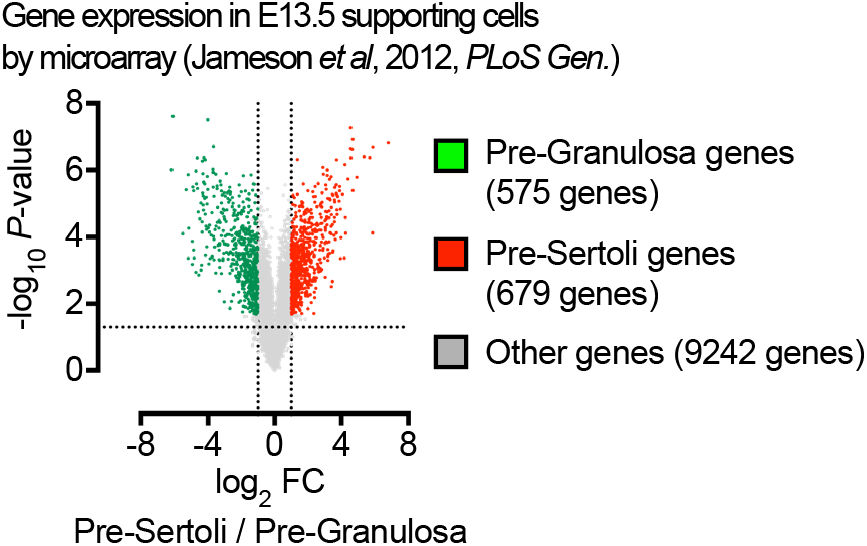
Expression profiles of specifically expressed genes in E13.5 granulosa cells. Microarray analysis of gene expression in E13.5 supporting cells (42). Genes with the criteria of 2-fold higher expression in E13.5 XX supporting cells and *P* < 0.05 were termed as “Pre-granulosa genes” and shown in green. Genes with the criteria of 2-fold higher expression in E13.5 XY supporting cells and *P* < 0.05 were termed as “Pre-Sertoli genes” and shown in red.

**Figure 5- figure supplement 2.**
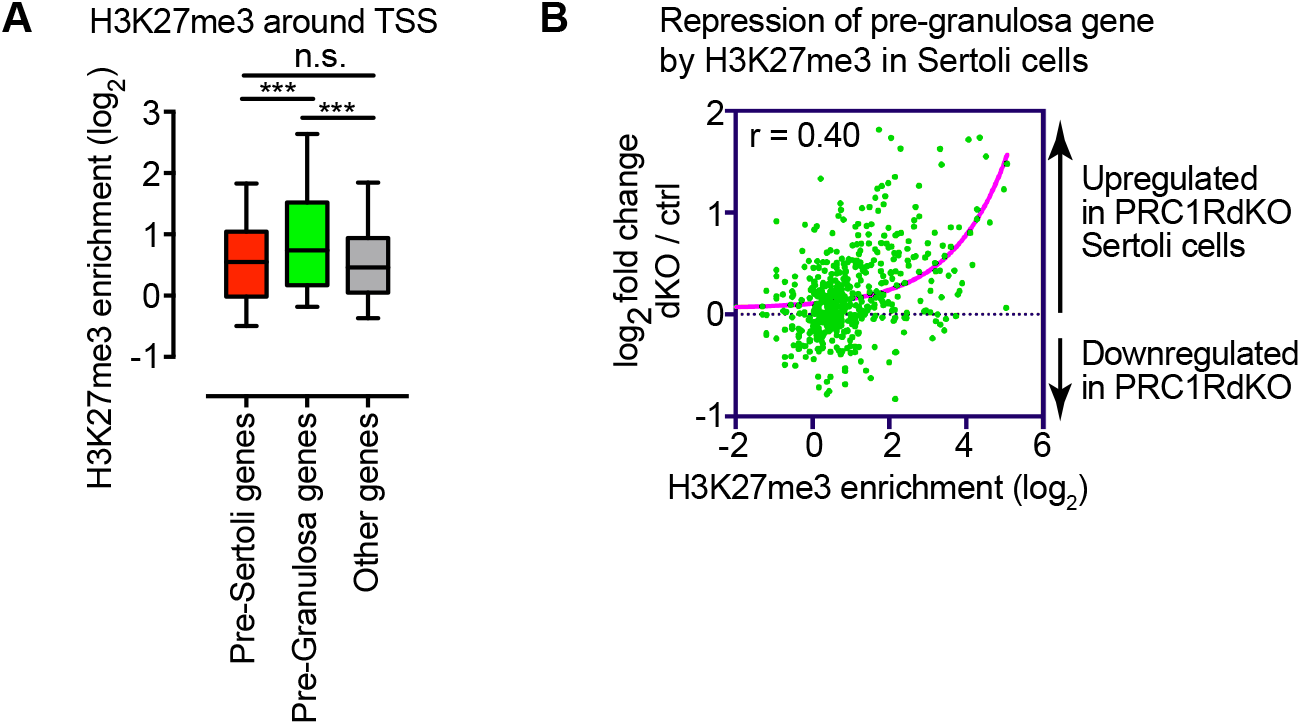
H3K27me3 is involved in the female gene regulatory network in Sertoli cells. (**A**)Box-and-whisker plots showing distributions of enrichment for H3K27me3 ChIP-seq data. Central bars represent medians, the boxes encompass 50% of the data points, and the whiskers indicate 90% of the data points. *\*\*\* P* < 0.0001, Mann-Whitney U tests. (**B**) Scatter plots showing the Pearson correlation between ChIP-seq enrichment (±2 kb around TSSs) and gene expression in Sertoli cells. A linear trendline is shown in blue.

